# Bioregionalization: From Wallace and Humboldt to deep-time paleoregion dynamics

**DOI:** 10.1101/2023.03.28.534545

**Authors:** Andrea Briega-Álvarez, Heike Siebert, Miguel Ángel Rodríguez, Sara Varela

## Abstract

Bioregionalization methods allow us to classify and map biogeographic units using data on species composition and traits. Here, we reviewed the evolution of the field during the last 70 years, seeking to summarize its history, and identify gaps and future avenues for research. Our results show that the aim of the studies using bioregionalization methods changed in time. First, bioregionalization were used to unveil the drivers of the observed spatial patterns of biodiversity on Earth, and to understand the role of dispersal limitations on the evolutionary history of clades, but recently, these methods are mostly used for conservation management. Further, data used to map biodiversity regions, the ones that we are now defining conservation strategies, are taxonomically and geographically biased, with a large percentage of the papers using vertebrate data from developed continents/countries. Finally, we show how key papers in the field, the ones with most citations, heavily depend on expert criteria and non-reproducible workflows, preventing direct comparison of maps of bioregions from different papers. Following our findings, we identified 3 gaps for the advance in the field, 1) We need to move beyond maps of vertebrate composition. Ideally, we need to increase the taxonomic diversity of the studies, but also to add other type of information, like data on species traits, genetic diversity, or phylogenetic distances. 2) we need reproducible and standardized methods 3) we need to further explore the temporal dimension of bioregions, to understand how they evolved through time.

## 1. Introduction

Bioregionalization methods allow us to cluster biodiversity and map regions with similar attributes (e.g. species composition) (Carstensen et al., 2013; Ebach, 2013; Ebach & Parenti, 2015; Ferrari, 2018; Mackey et al., 2008; McMahon et al., 2004; Morrone, 2014). These tools were used to investigate the role of historical events and current factors (e.g. plate tectonics, climate) on observed global biodiversity patterns (Bailey, 1983; Mackey et al., 2008; Mucina, 2019; Riddle & Hafner, 2010; Wallace, 1894). Further, bioregionalization methods were also used on international conservation programs (European Environment Agency, 2003; The Nature Conservancy, 2018; World Wildlife Fund, 2018), to map regions with similar biodiversity (e.g. Olson et al., 2001) and define management plans for them (Hobbs & McIntyre, 2005).

Different data has been used to identify spatial clusters of biodiversity. For example, Wallace and other seminal authors used species composition to identify historical units, or large biogeographic realms (Wallace, 1894). On the other hand, scientists such as Humboldt aimed to classify distinctive vegetation types independently of their species composition, based ecosystem physiognomy (Humboldt and Bonpland, 1807). The so-called biomes or vegetation types has been broadly used in biogeography and conservation biology. Thus, the initial use of regionalization tools was studying biogeography, how species composition might be used to unveil past dispersal constraints, and how the current distribution of species’ traits can be used to understand the role of climate in constraining species physiognomy. More recently, new subdivisions had been added, such as the ecoregions or zooregions, that divide the biogeographic realms into small geographic units with subclusters based on geography and species composition (Olson et al. 2007), while the initial goal of exploring theoretical questions about how life is distributed across space, moved towards a more practical use of these methods for conservation.

Research questions, data and approaches used by scientists working in ecology and evolution are school driven (Réale et al., 2020). In the case of regionalization, there are two main schools, a Taxonomic school that deals with classic biogeographical questions through taxonomic composition (e.g. the underlying drivers of species’ distributions), and a Functional school, more linked to conservation studies based on vegetation types (Mackey et al., 2008; Whittaker et al., 2013). Understanding the influence of such Schools in our research will give us clues to identify research biases and gaps in this field.

Here, we made a systematic review of regionalization papers of the last 70 years, and classified them in terms of data, methods and goals. With our analyses we seek to describe how bioregionalization studies have changed over time, as well as to identify gaps in the field, and future opportunities for research in biogeography.

## 2. Material and Methods

We built a bibliographic database on regionalization papers since 1951 until 2020 using three scientific repositories: Web of Science, Scopus and PubMed (see Supplementary Table 1). First, we split the bibliographic database into temporal periods of five years, counting the number of citations per period. Second, we identified the most cited papers during the study period, those that could have acted as milestones for the evolution of the field (see Table 1). Finally, we classified each article in our database according to six criteria, 1) methodology, 2) goals of the study, 3) type of biodiversity data, 4) type of evolutionary data, 5) major taxonomic group, and 6) geographic extent.

**Table 1.** Most cited papers of regionalization field considered as the ones before the stabilization of the curve of number of citations by year in decreasing order.

| Title | Cites<br>/year | Data | Goal | Method | Scale &<br>Location | Habitat | Type of<br>article |
| --- | --- | --- | --- | --- | --- | --- | --- |
| Olson, D. M. et al (2001). Terrestrial Ecoregions of the World: A New Map of Life on Earth. *BioScience*, 51(11), 933-938. | 214.1 | - | Biodiversity conservation | No numerical | Global | Terrestrial | Compilation |
| Spalding, M. D. et al (2007). Marine ecoregions of the world: a bioregionalization of coastal and shelf areas. *BioScience*, 57(7), 573-583. | 129.4 | - | Biodiversity conservation | No numerical | Global | Marine | Compilation |
| Holt, B. G. et al (2013). An update of Wallace's zoogeographic regions of the world. *Science*, 339(6115), 74-78. | 88.5 | Taxonomic & evolutionary | Methodological & establishing bioregions and potential drivers | (dis)similarity matrix-based decisions | Global | Terrestrial | Primary article |
| Morrone, J. J. (2014). Biogeographical regionalisation of the Neotropical region. *Zootaxa*, 3782(1), 1-110. | 84.9 | - | Description | No numerical | Neotropics | Terrestrial | Compilation |
| Abell, R. et al (2008). Freshwater ecoregions of the world: a new map of biogeographic units for freshwater biodiversity conservation. *BioScience*, 58(5), 403-414. | 80.1 | Taxonomic & evolutionary | Biodiversity conservation | No numerical | Global | Freshwater | Compilation |
| Omernik, J. M. (1987). Ecoregions of the conterminous United States. *Annals of the Association of American geographers*, 77(1), 118-125. | 54.1 | Functional | Biodiversity conservation | No numerical | USA | Terrestrial | Compilation |
| Morrone, J. J. (2006). Biogeographic areas and transition zones of Latin America and the Caribbean islands based on panbiogeographic and cladistic analyses of the entomofauna. *Annu. Rev. Entomol.*, 51, 467-494. | 46.5 | Taxonomic & evolutionary | Establishing bioregions and potential drivers | PAE & Panbiogeography | Latin America & Caribbean | Terrestrial | Primary article |
| Briggs, J. C., & Bowen, B. W. (2012). A realignment of marine biogeographic provinces with particular reference to fish distributions. *Journal of Biogeography*, 39(1), 12-30. | 42.8 | Taxonomic & evolutionary | Establishing bioregions and potential drivers | No numerical | Global | Marine | Compilation |

### 1. Methodology

We classified the articles according to the methodology used to regionalize in (dis)similarity matrix-based decisions; Cluster analysis; Network analyses; Original, ad-hoc numerical technique. We registered as “(dis)similarity matrix-based decisions” all procedures based on building a distance matrix (e.g. ordination analyses) except for clustering, a method that was classified in a separate category “Cluster analysis”. All graph-related classification approaches were classified as “Network analyses”. Because of the high number of other approaches to regionalize, each of them rarely employed, they were all registered under the category “Original, ad-hoc numerical technique” (see Supplementary Material for more details). Non-mathematical regionalizations (e.g. based on expert criteria, compilations) were classified as “No numerical” in this field (represented with the color white in Figure 2a).

**Figure 1.**
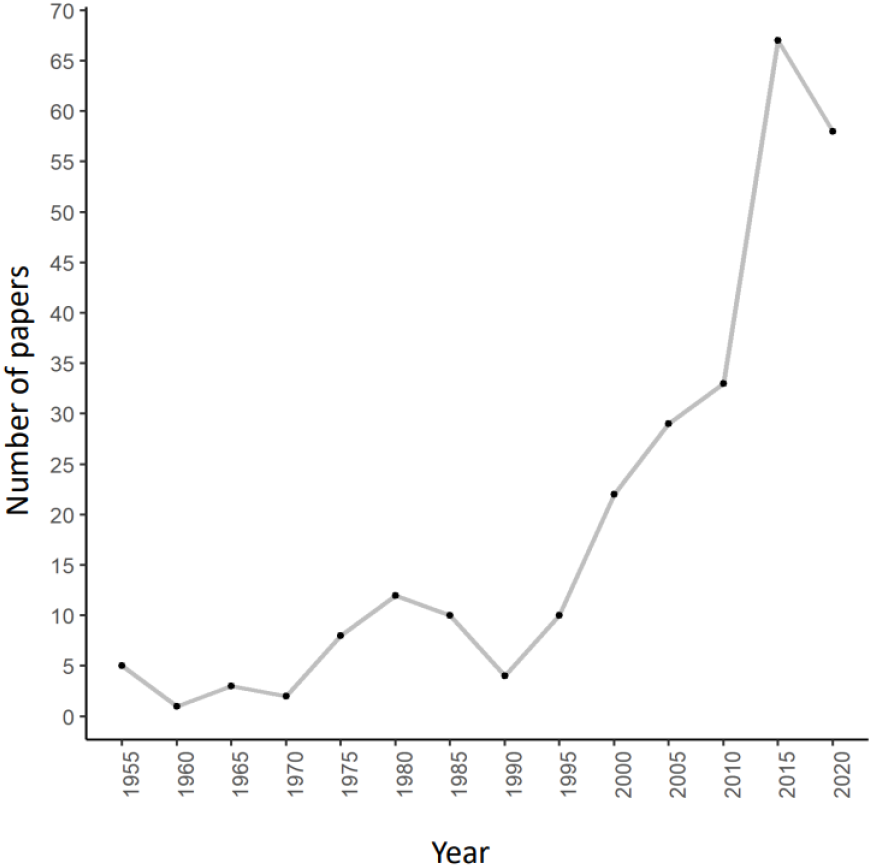
Number of regionalization articles each five years. The black circle represents the paper of Olson (2001) employed to split the database in two periods, marked in gray.

**Figure 2.**
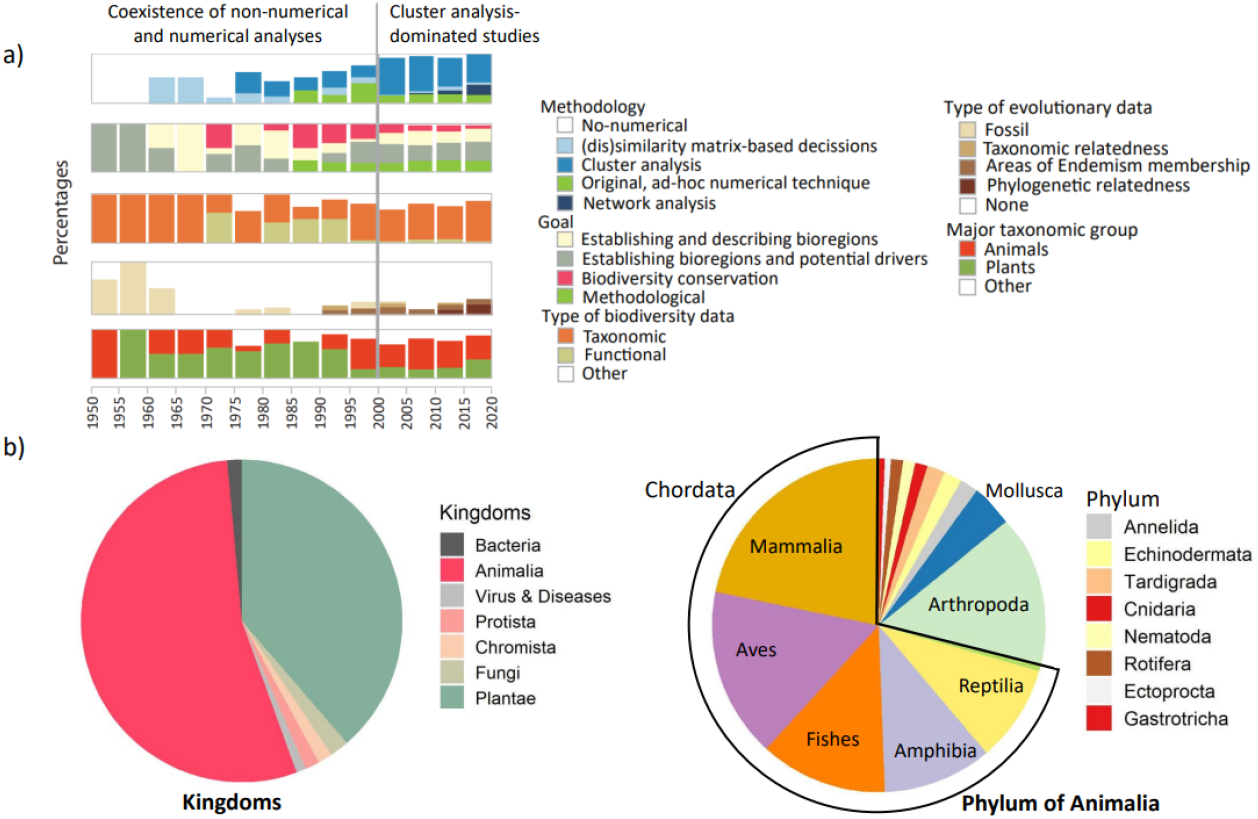
a) Proportion of articles that use each type of data, goal and methodology represented in barplots summarizing each 5 years in the database, with the “NA” category colored in white in each case. b) Proportion of phyla and classes for regionalized animals.

### 2. Goal

We classified the articles according to their main goal as establishing and describing bioregions; establishing bioregions and potential drivers; biodiversity conservation; methodological. The first two categories correspond to regionalizations whose main goal was to explore distribution patterns, either to just describe them or to better understand their underlying drivers (e.g. past events, climate). The other two include papers in which the main interest of the regions was their application either to nature conservation or environmental management (registered both as “biodiversity conservation”) or as an example to propose a new regionalization method or methodological improvement (classified as “methodological”).

### 3. Type of biodiversity data

We differentiated two biodiversity dimensions by classifying as “taxonomic” or “functional” regionalizations based on either taxonomic (e.g. species) or functional (e.g. vegetation types or physiognomic plant formations) characteristics, respectively. Complementarily, regionalizations that were no based on biodiversity characteristics, but on environmental information used as surrogates of them, were classified as “NA” (see proportions in white in Figure 2a for this and the additional “NA” categories presented below).

### 4. Type of evolutionary data

We used the category “taxonomical relatedness” to include regionalizations conducted on taxonomic data above the species level. These works generally assume that the hierarchy of taxonomical categories roughly reflects the timeline of the evolution (e.g. from ancient orders and families to recent species) and thus use upper taxonomic levels as a way to bring evolutionary information into the regionalization process. This follows the line of reasoning of A. R. Wallace, who argued that genera would be an adequate taxonomic level to reflect spatial assemblages whose onset predates the Pleistocene, while species would be too recent entities for that purpose (see Rueda et al. 2013 for a discussion of this topic). On the other hand, we used the category “Areas of endemism membership” to include regionalizations based on the identification of areas of endemism. These works assume that endemic taxa distributions are legacies of past events and thus carry important historical information. Besides, many of these methods, like recent versions of the so-called parsimony analysis of endemicity and the creation of areagrams, also include, at least partially, phylogenetic information (Ferrari, 2018). Additionally, we used the categories “fossil” and “phylogenetic relatedness” to comprise regionalizations based on fossil information and phylogenetic metrics (e.g. Holt et al., 2013), respectively. Finally, we assigned the category “NA” to papers not including any evolutionary information.

### 5. Major taxonomic group

Here, we split regionalizations based on animals and plants. Also, we kept the order, class and phylum used. Finally, we used the tag “NA” to reflect regionalizations based on other organisms or that employed environmental information only.

### 4. Geographical coverage (habitats, and continent or country of study)

We differentiated three habitat types –i.e., terrestrial, freshwater, and marine– and four geographical extents –i.e., global, continental, national, and regional (i.e., subnational)– noting for each sub-global regionalization the continent, country or region it was focusing on. We also built heatmaps at continental and national scales reflecting the number of regionalizations conducted at each scale (for visualization purposes, subnational-scale studies were treated as country level ones in this case, and freshwater and terrestrial regionalizations were combined). For this we used R coding (R Core Team, 2020) and the packages rworldmap, rgdal and RColorBrewer.

Finally, we performed Pearson Chi square tests by pairs of variables through STATISTICA V.14.0. to identify differences between the regionalization papers, as well as the relationships among them and with the goals and methods employed in the studies.

## 4. Results

We reviewed 233 papers from the period 1950-2020, of which the most cited one was “Terrestrial ecoregions of the World” by D. M. Olson et al. (2001). Indeed, with 4282 citations in total, and an average of 214.1 citations per year (Table 1), this paper nearly doubled the citation rate of the one scoring second (Spalding et al., 2007), and was published right when the field was acquiring momentum in terms of the number of papers being published every year (see Figure 1). Interestingly, although Olson et al. (2001) presented a non-numerical, global scale synthesis of previous regionalizations aimed at being the basis for conservation studies and initiatives, it was published in a year that marked the transition to a period in which numerical methods, particularly cluster analysis-based ones, became virtually the norm in bioregionalization studies (Figure 2a). Thus, we split our database into two major periods: namely 1950-2000 and 2001-2020, comprising 64 and 169 papers each (see Figure 2a).

Regarding the study goals, they passed from defining bioregions to start generating bioregions for conservation purposes in the 70s, or producing methodological studies since the mid-80s. The early 2000s also marked the transition to a period were the objective of establishing regions and their (historical, ecological, and geographical) drivers rated first, and producing regions for conservation purposes rated last. Other aspects, like merely defining bioregions and describing them verbally or focusing on methodological aspects had intermediate interest (see Figure 2a).

As for the type of biodiversity data used, taxa, and particularly species distributions, were used in 97% of the cases. Functional data was also important from 1970s to 1990s, most of the second half of the first period (see Figure 2a).

Regarding evolution, except for the earliest 15 years in which fossils were often used to incorporate past information into bioregionalizations, the norm has been to speculate rather than using data-based analyses. Still, different evolutionary data sources have been employed in some studies, including not only fossils, but also data on areas of endemism, taxonomic relatedness, and, in the last ten years, on phylogenetic relatedness (see Figure 2a), likely due to the strong influence exerted by the highly cited paper by Holt et al. published in 2013 (see Table 1).

Finally, the earliest period (before 2001) was focused firstly on plants and then on animals, while studies devoted to other organisms were scarce, and only appeared since the mid-70s. In contrast, the most recent period was clearly dominated by studies focusing on animals, followed by those using either plants or other organisms, in similar proportions (see Figure 2a).

Regarding spatial extents and habitats, study proportions did not differ much before and after 2001, nor across geographical scales, with terrestrial habitats clearly dominating the field (ranging from 81.3% to 73.3% of the papers across periods and extents), followed by marine and freshwater habitats (Table 2). At the continent and country levels, North America and USA scored in the first place (14 before 2001 in each case, and 13 and 6 respectively afterwards). At this time, South America, Australia and Europe joined that trend with 9 articles each and China overcame the regionalization level of USA (7 articles). Considering all database, the most regionalized places have been North and South America, Europe, USA, Australia and China. Regarding marine regions, at the continental/oceanic level they were spread over the world, while at the national level they were highly concentrated in the Mediterranean Sea (Figure 3).

**Table 2.** Percentages of regionalizations by period and each of the classified variables and their categories.

| **Categories** | **Variables** | **1950-**<br>**2000** | **2001-**<br>**2020** |
| --- | --- | --- | --- |
| Methodology | Cluster analysis | 22.9 | 57.4 |
|  | (dis)similarity matrix-based decisions | 14.3 | 5.1 |
|  | Network analyses | 0 | 10.2 |
|  | Original, ad-hoc numerical technique | 14.3 | 16.5 |
|  | No numerical | 48.6 | 10.8 |
| Goal | Establishing bioregions and potential drivers | 39.1 | 37.1 |
|  | Establishing and describing bioregions | 25.0 | 25.3 |
|  | Biodiversity conservation | 25.0 | 11.8 |
|  | Methodological | 10.9 | 22.9 |
| Type of biodiversity data | Taxonomic | 62.5 | 74.6 |
|  | (Species only) | (60.9) | (72.3) |
|  | Functional | 25.0 | 5.9 |
|  | (Vegetation forms only) | (18.1) | (4.4) |
| Type of evolutionary data | Fossil | 13.9 | 1.3 |
|  | Taxonomical relatedness | 1.5 | 2.5 |
|  | Areas of endemism membership | 4.6 | 11.3 |
|  | Phylogenetic relatedness | 0 | 6.9 |
| Major taxonomic group | Animals | 40.6 | 54.1 |
|  | Plants (terrestrial plus aquatic) | 46.9 | 25.5 |
|  | (terrestrial only) | (37.5) | (24.9) |
| Geographical scale | Global/terrestrial | 21.9 | 13.7 |
|  | Global/freshwater | 1.6 | 1.2 |
|  | Global/marine | 3.1 | 5.0 |
|  | Continental/terrestrial | 26.6 | 19.9 |
|  | Continental/freshwater | 1.6 | 1.9 |
|  | Continental/marine | 6.3 | 4.3 |
|  | National/terrestrial | 26.6 | 26.7 |
|  | National/freshwater | 1.6 | 1.9 |
|  | National/marine | 4.7 | 6.2 |
|  | Regional/terrestrial | 7.8 | 13.7 |
|  | Regional/freshwater | 0 | 5.0 |
|  | Regional/marine | 1.6 | 1.9 |

**Figure 3.**
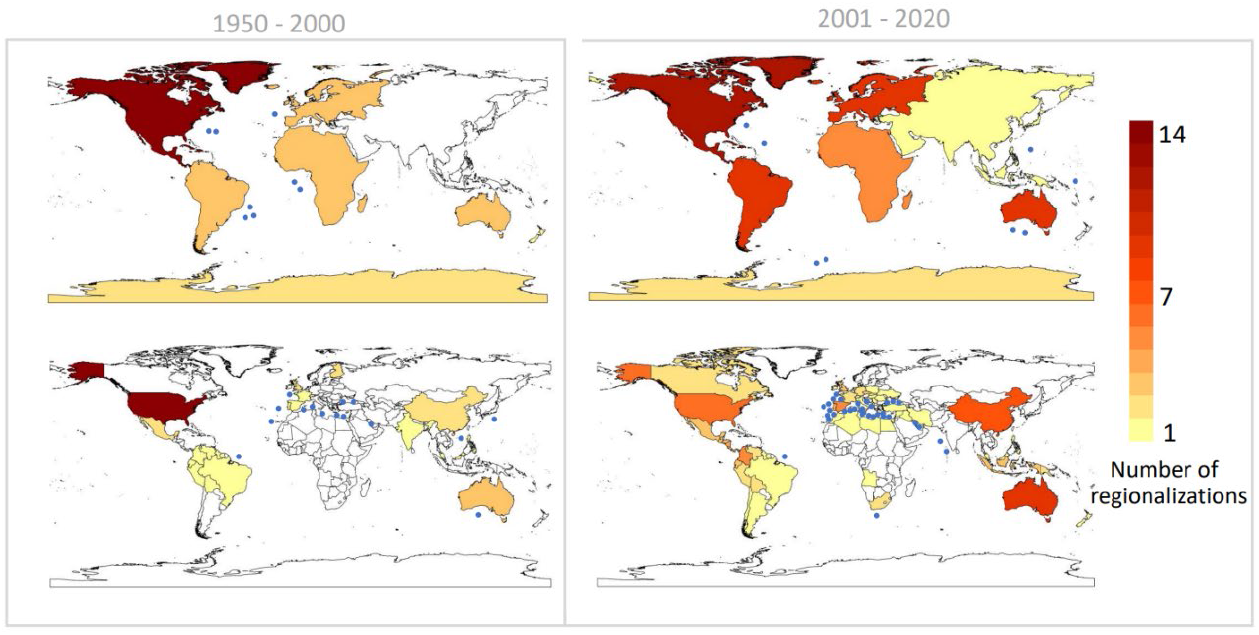
Heatmaps of number of regionalizations by continent and country for each period and proportions of regionalizations by geographical scale and period. Since the boundaries for marine regionalizations were very variable among articles, they are marked as blue circles for visualization purposes.

Finally, we identify two rough groups of articles mirroring the classic Taxonomic and Functional schools of Biogeography. 1) The Taxonomic group encloses most articles and is characterized by the dominant use of taxonomic over functional data (X^2^ (1, N = 191) = 23.08, p < .001, supp. Table 2j), mainly based on species (Table 2). The use of species is also significatively related to articles with no-conservation goals (X^2^ (3, N = 187) = 61.93, p < .001, supp. Table 2c). 2) The Functional group is based on functional data, mainly using vegetation types (supp. Table 2j) in papers with conservation goals (supp. Table 2c) and through no numerical approaches (X^2^ (3, N = 147) = 40.10, p < .001, supp. Table 2h). This group (altogether, the use of functional data, the conservation goal and no numerical approaches) was more popular before 2001 (Table 2).

In both groups, the evolutionary dimension was mostly unexplored, but its rare consideration occurs significatively more in basic bioregionalization articles than in the applied conservationists or methodological ones (X^2^ (3, N = 228) = 22.80, p < .001, supp. Table 2a).

Regarding the most impactful articles in the last 70 years of regionalization field, we selected 9 (supplementary Figure 1). Most of them are classic examples of the Functional group, conservation articles based on vegetation types-except for (Abell et al., 2008)-applying no numerical methods and/or compilations of previous regions to get large scale regionalizations (Table 1). In this shortlist, we also have an article that has a key impact in how researchers are including evolutionary data in bioregionalization, a paper that suggests the direct inclusion of phylogenetic metrics, particularly as phylogenetic distances among geographic cells, to delineate bioregions through clustering algorithms (Holt et al., 2013).

## 5. Discussion

Here, we have classified 233 articles in relation to their goals, data, and methods (Figure 1 and 2). Our results confirm the breach between Taxonomic and Functional schools (Figure 2), already noticed by other researchers (Mackey et al., 2008; Whittaker et al., 2013). However, despite the clear dominance of the Taxonomic school (by number), there is a mismatch between impactful and “common” regionalization articles. The papers with the greatest impact are studies using functional categories for conservation (see Table 1). The Functional school consists mainly in conservation articles were regions are based on vegetation types and no numerical approaches, such as compilations and/or expert criteria. Although many of these articles intend to delineate historically and ecologically meaningful regions (Olson et al., 2001; Udvardy, 1975), the lack of standardized data forced them to use the available information in a no-reproducible way.

Interestingly, there are also papers with a high impact, that used mathematical approaches to build their regions. For instance, the third most impactful paper (Table 1) added phylogenetic relationships into the regionalization process (Holt, 2013). The phylogeny is included as phylogenetic similarity among geographic cells (phylo-beta-sim) in the calculation of the distance matrix, a method vastly applied in the last decade (Daru et al., 2016, 2017; Hattab et al., 2015; Li et al., 2015; Pataro et al., 2020; Slik et al., 2018; Ye et al., 2019). Several authors critic that phylo-beta-sim fails in grouping in the same region clades with different evolutionary histories (Kreft & Jetz, 2013), a problem that has been solved in a posterior refinement of the method through the use of “fuzzy” sets (Maestri & Duarte, 2020).

### Biases in biodiversity data

Regionalization, such as other fields of biology, suffer from the use of biased/incomplete sets of data (Figure 2). Vertebrates (particularly mammals and birds) are strongly overrepresented, while plants, fungi and invertebrates (specially insects) are underrepresented (Scholl & Wiens, 2016). The bias towards animals over plants is more pronounced in this field than in other areas of biogeography (Bonnet et al., 2002; Donaldson et al., 2017), especially after 2001, with an exponential growth in number of articles using vertebrates. This could be explained by a combination of research interest towards vertebrates, together with the greater and sooner availability of distribution data for vertebrate species (IUCN, 2008). However, the bias towards birds and mammals (19.5 and 14.7% of the articles respectively), is not so high as in other ecological studies, with a 44 and 27% of the papers based on them (Bonnet et al., 2002). A possible reason could be that the databases on distribution on reptiles and amphibians were not available at the time when other ecological studies were made (IUCN, 2008). Besides, despite birds have been traditionally more sampled (Hughes et al., 2020), mammals have been more regionalized, probably because this was the first group for which global distribution data was available (IUCN, 2008). On the other hand, although invertebrates are underrepresented, they are more studied through regionalization (36.3% of the articles) than in conservation biology (31.92%, see Donaldson et al., 2017). Most invertebrate regionalizations were based on insects, followed by crustaceans (see Figure 2b), like the delineation of global marine regions for conservation (Costello et al., 2017).

In general, our results show that, although bioregions were first made to describe and synthesize species distribution patterns, their uses have diversified along time. The Big Data revolution of the 21^st^ century has been fueling bioregionalization research. Notably, the early 2000s also witnessed the raising of evolutionary ecology (Réale et al., 2020), and the blooming of global biodiversity databases (IUCN, 2008), two aspects that, as we shall see, were highly influential on the data, methods and goals characterizing modern bioregionalizations. We detected that bioregionalization studies follow the same trend than other studies in biology, were data availability is still deeply biased (Hughes et al., 2020).

Finally, our data indicate that information on evolution (e.g. phylogenies) is less included in articles dealing with conservation than in articles with other goals. Evolutionary uniqueness has been shown to be a key conservation aspect (Isaac et al., 2012), but not only because of the intrinsic value of preserving evolutionary unique species (Rosauer et al., 2009), but also because its relation to the susceptibility of species to extinction (Fritz & Purvis, 2010). Thus, we believe that regionalization analysis with conservation goals would benefit if including phylogenetic information.

### Geographic biases

Besides, bioregionalization research, as other biodiversity studies, is also affected by geographical biases (see Figure 3), both among habitats (terrestrial vs. aquatic) and across continents. In general, terrestrial habitats have greater sampling and then, more data, than aquatic environments (Hughes et al., 2020). Regarding the geographic biases in the papers dealing with marine data, the Mediterranean Sea is the most regionalized sea/ocean. A possible explanation could be its small size, since the availability of Biodiversity data in the marine habitats seems to be related to the distance to the coastline (Hughes et al., 2020), and that it is part of European countries with high GDP and therefore high scientific investment.

The number of regionalizations by country is related to the economic development (simple correlation between the Gross Domestic Product (International Monetary Fund, 2021) and number of regionalizations: r=0.835; p-value << 0.001). North America and United States were the first continent and country in being repeatedly regionalized, probably because of its more recent economic and scientific development than other countries, together with the old history of biological conservation of North America, where ecoregions were first defined and delineated with conservation purposes (Bailey & Hogg, 1986). Generally, research efforts are focused on developed countries (Hughes et al., 2020; Silva et al., 2020). For example, an 82% of distribution records in GBIF and OBIS databases are from only a 4% of countries of the world, particularly from US, Europe, Australia and South Africa (Hughes et al., 2020) while countries with less accessible habitats and small countries are understudied (Pyšek et al., 2008; Silva et al., 2020; Trimble & Aarde, 2012).

### Future avenues for research

Bioregionalization methods can be used to understand how life evolved and distributed across space and time, and to explore how and when commonly used variables, such as climate, triggered biogeographic changes. Bioregionalization is considered a very diverse and even conceptually conflicted field (Ferrari, 2018; McMahon et al., 2004), including many different definitions of biogeographical regions and approaches to define them (Carstensen et al., 2013b; Ebach & Parenti, 2015; Mackey et al., 2008), but in this review we found out that it is quite uniform, especially after 2001 (Figure 2a, Table 2). In this sense, the paper of Kreft and Jetz (2010) could have played a central role influencing the field onwards. Rather than its original evolutionary meaning (Wallace, 1876, 1894), bioregions are currently used to map species pools (gamma diversity).

However, recently, paleontologists are starting to use regionalization tools to map deep time biodiversity changes (Kocsis et al. 2021). New paleoclimatic reconstructions on past climate is allowing us to start exploring how and to which extent, climate might be related to the evolution and extinction of life on Earth. Thus, we believe that as a consequence of the expansion of the field towards paleontological reconstructions, we might see a revolution in the field of biogeography. New avenues for research will include the combination of fossil data with current data on species occurrences, to understand long term dynamics on bioregions, and how different factors (e.g. climate change and plate tectonics) played a role on the emergence of distinct evolutionary regions.

Following our findings, we identified 3 future avenues for research for further exploring the spatial clustering of life composition and attributes, 1) moving beyond maps of vertebrate composition: we need taxonomically diverse data, and data on species traits, genetic diversity, phylogenetic distances. 2) moving towards a more reproducible and standardized procedures: we need to compare map results from different papers. 3) adding the temporal dimension: we need to understand how bioregions evolved through time.

## Supporting information

Briega_et_al_Sup_Material

## Author contributions

All authors contributed to design the study. ABA collected data, performed analyses and wrote the first draft of the manuscript. SV, HS, MAR contributed with major revisions to the final manuscript.

