## Supplementary material for "Bioregionalization: From Wallace and Humboldt to deep-time paleoregion dynamics": Briega_et_al_Sup_Material

### Supplementary information

#### Further explanations for the classification of the database

39 articles are classified as using a data type "Not-applicable". 23 of them classify environmental/physical variables only, 14 are compilations of previous regionalization studies and 2 are compilations that also consider such environmental/physical variables. All these papers are also classified as "Not-applicable" taxonomical categories, plus two more that use different taxa than animals and plants, particularly fungi and human diseases respectively.

5 articles are classified as including evolutionary history in the regionalization but with "other" approaches than the ones included in table 2. Such approaches consist in expert criteria, references to the landscape in the glacial periods, tectonic movements and references to the "geoelement" concept, that is historical by definition (Udvardy, 1975).

5 papers are not classified into the goals of table 2 because their main goal was not directly related with the regionalization exercise. The delineation of regions in that cases was applied as additional support to answer different questions, e.g. if a species complex could be considered as different species or not.

41 papers are classified as "others" in method since they employ a different mathematical method than the ones included in table 2. That methods, very rarely used (only 1 or 2 articles use each of them) except for the methods to identify areas of endemism, are listed below:

- Isocluster
- Classification and regression trees (CART)
- Bayesian tree models and linear regressions

- Fitch’s parsimony method
- Barrier Analysis
- Logistic discrimination (LGD) and linear discriminant functions (LDF)
- Mixture of experts model framework, which includes environmental and biological variables but also expert criteria.
- Methods for the identification of areas of endemism (Parsimony Analysis of Endemicity -PAE-, Analysis of Endemicity -NDM and VNDM-, etc)
- Statistical matrices
- Panbiogeography
- Spatial clustering methods
- GIS related methods
- GADM models

Supplementary Table 1. Advanced search in WOS format.

| | |
|---|---|
| (TI=bioregions OR<br>TI=biorregions OR<br>TI=phyloregions OR<br><br>TI="biological regions" OR<br>TI="biogeographic regions" OR<br>TI="biogeographical regions" OR<br>TI="phytogeographical regions" OR<br>TI="phytogeographic regions" OR<br>TI="floral regions" OR<br>TI="floristic regions" OR<br>TI="zoogeographic regions" OR<br>TI="zoogeographical regions" OR<br>TI="phylogenetic regions" OR<br>TI="zooregion" OR<br>TI="phytoregion" OR<br><br>TI="phytogeographical zone" OR<br><br>TI="biological provinces" OR<br>TI="biogeographic provinces" OR<br>TI="biogeographical provinces" OR<br>TI="phytogeographical provinces" OR<br>TI="phytogeographic provinces" OR<br>TI="floral provinces" OR<br>TI="floristic provinces" OR<br>TI="zoogeographic provinces" OR | TI="floristic provincialization" OR<br>TI="zoogeographic provincialization" OR<br>TI="zoogeographical provincialization" OR<br>TI="phylogenetic provincialization" OR<br><br>TI="biogeographic structure" OR<br>TI="biogeographical structure" OR<br>TI="phytogeographical structure" OR<br>TI="phytogeographic structure" OR<br>TI="zoogeographic structure" OR<br>TI="zoogeographical structure" OR<br><br>TI="biological units" OR<br>TI="biogeographic units" OR<br>TI="biogeographical units" OR<br>TI="phytogeographical units" OR<br>TI="phytogeographic units" OR<br>TI="floral units" OR<br>TI="floristic units" OR<br>TI="zoogeographic units" OR<br>TI="zoogeographical units" OR<br><br>TI=bioregionalisation OR<br>TI=bioregionalization OR |

| | |
|---|---|
| TI="zoogeographical provinces" OR<br>TI="phylogenetic provinces" OR<br><br>TI="floral kingdoms" OR<br><br>TI="biological regionalization" OR<br>TI="biogeographic regionalization" OR<br>TI="biogeographical regionalization" OR<br>TI="phytogeographical regionalization" OR<br>TI="phytogeographic regionalization" OR<br>TI="floral regionalization" OR<br>TI="floristic regionalization" OR<br>TI="zoogeographic regionalization" OR<br>TI="zoogeographical regionalization" OR<br>TI="phylogenetic regionalization" OR<br><br>TI="biological regionalisation" OR<br>TI="biogeographic regionalisation" OR<br>TI="biogeographical regionalisation" OR<br>TI="phytogeographical regionalisation" OR<br>TI="phytogeographic regionalisation" OR<br>TI="floral regionalisation" OR<br>TI="floristic regionalisation" OR<br>TI="zoogeographic regionalisation" OR<br>TI="zoogeographical regionalisation" OR<br>TI="phylogenetic regionalisation" OR<br><br>TI="biological provincialization" OR<br>TI="biogeographic provincialization" OR<br>TI="biogeographical provincialization" OR<br>TI="phytogeographical provincialization" OR<br>TI="phytogeographic provincialization" OR<br>TI="floral provincialization" OR | TI="biogeographic provinces" OR<br>TI="biogeographical provinces"<br><br>TI="biological communities" OR<br>TI="ecological communities" OR<br>TI="biogeographic communities" OR<br>TI="biogeographical communities" OR<br>TI="phytogeographical communities" OR<br>TI="phytogeographic communities" OR<br>TI="floral communities" OR<br>TI="floristic communities" OR<br>TI="zoogeographic communities" OR<br>TI="zoogeographical communities" OR<br>TI="phylogenetic communities"<br><br>TI="ecoregions" OR<br>TI="ecological regions" OR<br>TI="ecoregionalization" OR<br>TI="eco-regionalization" OR<br>TI="conservation regions" OR<br>TI="regions for conservation") AND<br><br>(SU="Biodiversity & Conservation" OR<br>SU="Environmental Sciences & Ecology" OR<br>SU="Evolutionary Biology" OR<br>SU="Marine & Freshwater Biology" OR<br>SU="Mycology" OR<br>SU="Palentology" OR<br>SU="Plant sciences" OR<br>SU="Zoology" OR<br>SU="Oceanography") |

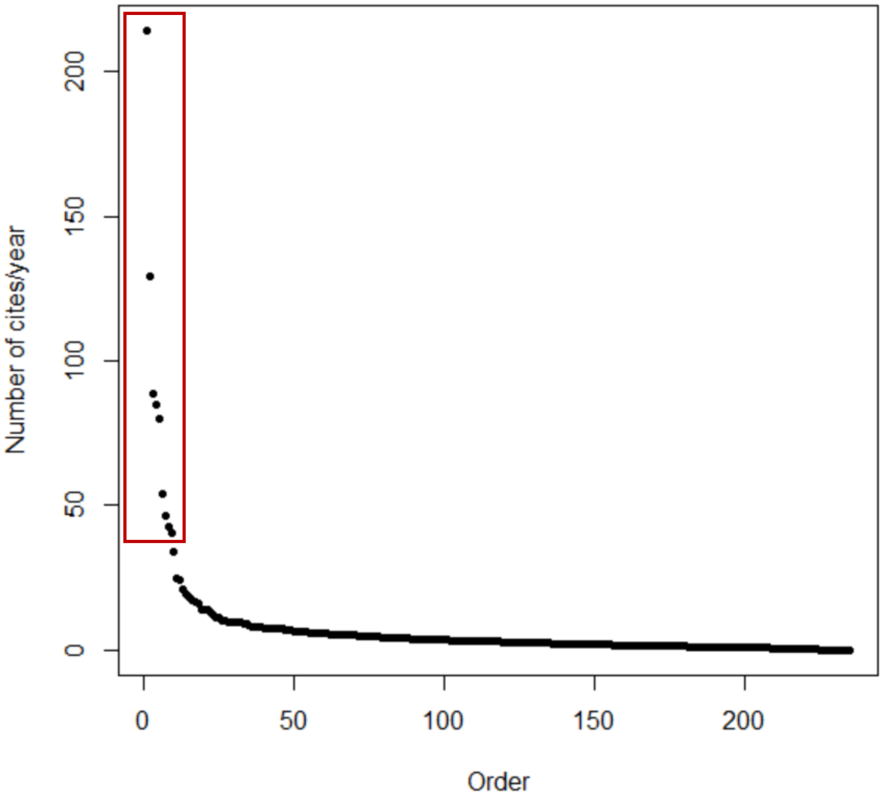

Supplementary figure 1. Number of citations in Web of Science by year for the bibliographic database. The red square highlights the most cited articles as the previous ones to the stabilization of the curve, detailed in Table 1.

Supplementary tables 2 a-j. Contingency tables for Chi square test. Significative biases are marked in bold and marginally significative are underlined.

Supplementary table 2a. Contingency table of type of the inclusion of evolutionary data vs goal.

| | | Inclusion of evolutionary data | | |
|---|---|---|---|---|
| | | Included | Not included | Total |
| Goal | Establishing bioregions and potential drivers | **36** | **52** | 88 |
| | Establishing and describing bioregions | **11** | **48** | 59 |
| | Biodiversity conservation | 2 | **34** | 36 |
| | Methodological | 7 | **38** | 45 |
| | Total | 56 | 172 | 228 |

Supplementary table 2b. Contingency table of methodology vs goal.

| | | Methodology | | | | |
|---|---|---|---|---|---|---|
| | | Cluster analysis | (dis)similarity matrix-based decisions | Network analyses | No numerical | Total |
| Goal | Establishing bioregions and potential drivers | **45** | 3 | 3 | **19** | 70 |
| | Establishing and describing bioregions | **24** | 5 | 2 | **13** | 44 |
| | Biodiversity conservation | **12** | 2 | 0 | **20** | 34 |
| | Methodological | **21** | 1 | 4 | 2 | 28 |
| | Total | 102 | 11 | 9 | 54 | 176 |

Supplementary table 2c. Contingency table of type of biodiversity data vs goal.

| | | Type of biodiversity data | | |
|---|---|---|---|---|
| | | **Taxonomic** | **Functional** | **Total** |
| **Goal** | **Establishing bioregions and potential drivers** | **80** | 2 | 82 |
| | **Establishing and describing bioregions** | **43** | 6 | 20 |
| | **Biodiversity conservation** | 6 | **14** | 82 |
| | **Methodological** | **32** | 4 | 36 |
| | **Total** | 161 | 26 | 187 |

Supplementary table 2d. Contingency table of major taxonomic group vs goal.

| | | Major taxonomic group | | |
|---|---|---|---|---|
| | | **Animals** | **Plants** | **Total** |
| **Goal** | **Establishing bioregions and potential drivers** | 49 | 32 | 81 |
| | **Establishing and describing bioregions** | 28 | 21 | 49 |
| | **Biodiversity conservation** | 9 | 11 | 20 |
| | **Methodological** | 28 | 8 | 36 |
| | **Total** | 114 | 72 | 186 |

Supplementary table 2e. Contingency table of methodology vs inclusion of evolutionary data

| | | Methodology | | | | |
|---|---|---|---|---|---|---|
| | | **Cluster analysis** | **(dis)similarity matrix-based decisions** | **Network analyses** | **No numerical** | **Total** |
| **Inclusion of evolutionary data** | **Included** | 18 | 0 | 10 | 14 | 34 |
| | **Not included** | 84 | 11 | 2 | 40 | 145 |
| | **Total** | 102 | 11 | 12 | 54 | 179 |

Supplementary table 2f. Contingency table of type of biodiversity data vs inclusion of evolutionary data.

| | | **Type of biodiversity data** | | |
|---|---|---|---|---|
| | | **Taxonomic** | **Functional** | **Total** |
| **Inclusion of evolutionary data** | **Included** | 53 | 2 | 55 |
| | **Not included** | 113 | 24 | 137 |
| | **Total** | 166 | 26 | 192 |

Supplementary table 2g. Contingency table of major taxonomic group vs inclusion of evolutionary data.

| | | **Major taxonomic group** | | |
|---|---|---|---|---|
| | | **Animals** | **Plants** | **Total** |
| **Inclusion of evolutionary data** | **Included** | 34 | 20 | 54 |
| | **Not included** | 84 | 53 | 137 |
| | **Total** | 118 | 73 | 191 |

Supplementary table 2h. Contingency table of type of biodiversity data vs methodology.

| | | **Type of biodiversity data** | | |
|---|---|---|---|---|
| | | **Taxonomic** | **Functional** | **Total** |
| **Methodology** | **Cluster analysis** | **86** | 6 | 92 |
| | **(dis)similarity matrix-based decisions** | 9 | 0 | 9 |
| | **Network analyses** | **12** | 0 | 12 |
| | **No numerical** | **17** | **17** | 34 |
| | **Total** | 124 | 23 | 147 |

Supplementary table 2i. Contingency table of major taxonomic group vs methodology.

| | | **Major taxonomic group** | | |
|---|---|---|---|---|
| | | **Animals** | **Plants** | **Total** |
| **Methods** | **Clustering algorithms** | **57** | **35** | 92 |
| | **Distance methods** | 9 | 0 | 9 |
| | **Graph analysis** | **11** | 1 | 12 |
| | **No mathematical** | 10 | **24** | 34 |
| | **Total** | 87 | 60 | 147 |

Supplementary table 2j. Contingency table of major taxonomic group vs type of biodiversity data.

| | | Major taxonomic group | | |
|---|---|---|---|---|
| | | **Animals** | **Plants** | **Total** |
| **Type of biodiversity data** | **Taxonomic** | **113** | **52** | 165 |
| | **Functional** | 5 | **21** | 26 |
| | **Total** | 73 | 118 | 191 |
